# Thymosinβ4 enhances liver regeneration

**DOI:** 10.64898/2026.06.22.733089

**Authors:** Li Xiankui

## Abstract

Thymosin β4 (Tβ4) is a conserved acidic polypeptide with 43-amino acids participating in multiple pathophysiological processes. In this study *in vivo* effects of Tβ4 on liver regeneration are investigated in carbon-tetrachloride (CCL4) induced rodent animal liver jury models. Results illustrate that exogenous Tβ4 treatment significantly reduced CCL4-rendered liver necrosis around central vein. At 48 hours after CCL4 insults hepatocytes proliferation occur mainly around the periportal area, while hepatocytes proliferation around the necrosis area is prominently increased by exogenous Tβ4 treatment. The holistic proliferation level of liver tissues are also enhanced by exogenous Tβ4. Hepatocyte proliferation activities negatively correlate with the necrosis extent of the liver tissue. These results suggested firstly exogenous Tβ4 treatment could enhance liver regeneration and exhibit prosperous potential for application in clinical conditions such as liver transplantation.

## Introduction

Thymosin β4 (Tβ4) is a highly conserved 43-amino acid polypeptide that exhibits multiple biological functions, including anti-inflammatory, angiogenic, and tissue reparative activities^1^. Recent studies indicates Tβ4 is a therapeutic regenerative peptide beneficial to tissue regeneration^2^. Tβ4 have been shown to extend the time window of neonatal mouse heart regeneration and promote cardiomyocyte proliferation^3 4^, accelerated re-epithelialization during cutaneous wound healing^5^. Tβ4 ophthalmic solution has entered Phase III clinical trials for neurotrophic keratopathy, showing favorable effects on corneal epithelial repair^6^. At present, effects of Tβ4 on liver regeneration has not been investigated and liver regenerative capacity is a major determinant liver failure^7^. In this study liver regeneration after carbon-tetrachloride (CCL4) insults is investigated in a murine liver injury model and results indicate that exogenous Tβ4 treatment enhances hepatocytes proliferation and reduces CCL4-rendered liver necrosis.

## Methods

### Statement of Ethics

Animal experiments were approved by local Ethics Committee for Animal Care and Use at Tianjin Medical University (Ethic No. TMUaMEC2015003). All animals were treated humanely in accordance with Guide for the Care and Use of Laboratory Animals (Institute of Laboratory Animal Resource, 1996, Nat. Acad. Press).

### Murine liver injury model

Six-weeks old male Balb/c mice (∼20 g) were purchased from the Branch of National Breeder Center of Rodents (Beijing) and raised in specific-pathogen-free (SPF) facilities (Experimental Animal Center at Tianjin Medical University). 12-h light/dark cycle and *ad libitum* access to food and water were offered. Mice were acclimatized for 7 days in the SPF environment and then used for the following experiments. Pentobarbital was used to anesthetize mice and euthanasia was performed at the end of the experiments.

Mice were divided into two groups (n=5), CCl4 treatment group and CCL4+Tβ4 treatment group. In CCl4 group, mice received a single intraperitoneal injection of 0.2 ml/kg.bw (bw, body weight) CCl4 in olive oil to induce liver injury and some doses of saline injection. The saline volume of every dose equal the volume of the same volume of Tβ4 in CCL4+Tβ4 group. In CCL4+Tβ4 group besides one dose of CCL4 injection as CCL4 group mice also received intraperitoneal Tβ4 administration at 0 hour, 2 hours, 4 hours, 6 hours after CCl4 injection. Tβ4 used in these experiments was bought from GL Biochem. (Shanghai) Ltd. The purity of Tβ4 was higher than 98% as determined by HPLC. During experiments Tβ4 was dissolved in 0.9% saline for injection.

### Histology

After sacrificing the mice liver tissues were collected. 10% neutral buffered formalin was used to fix the liver tissues followed by routinely processing. After paraffin embedding the liver tissues were cut into 4-μm thick sections. And the tissue slides were stained with standard hematoxylin-eosin (H.E) staining. Images were captured on 3DHISTECH digital pathology system. Images with 40x magnification were analyzed for necrosis area quantification using free software QuPath (https://qupath.github.io/) and expressed as proportion in percentage. Necrosis area were recognized by automatic training procedures in QuPath and then all the images were processed in batch.

### Immunohistochemistry (IHC)

Liver sections were subject to deparaffinization, complete rehydration and 3% H2O2 for 10 min blocking endogenous peroxidase activities. Antigen retrieval was done in citrate buffer (pH=6.0) in a microwave oven for 15min. 5% BSA was used to block non-specific protein binding. Then the slides were incubated with primary anti-PCNA antibody (PC10, 1:200, eBioscience) overnight at 4 °C in a humidified chamber. After twice washing with PBS the liver slides were sequentially probed and amplified with a biotinylated secondary antibody and streptavidin–avidin–peroxidase complex. Color was developed with diaminobenzidine (DAB) Chromogen. Finally, slides were lightly counter-stained with hematoxylin and mounted with mounting medium.

For IHC, all the liver slides were stained in parallel using identical staining conditions. After IHC the whole side was scanned on 3DHISTECH digital pathology system and images were viewed and captured using Pannoromic Viewer software. Free software QuPath (https://qupath.github.io/) was using to recognize the PNCA-positive cells in 3 random fields (200x magnification). All the images were analyzed under the same parameters. The staining intensity and PCNA-positive area proportion were semi-quantified and expressed as comprehensive H-score^8^. Nuclear H-score and cytoplasmic H-score were measured separately in order separately evaluate the PCNA levels in nucleus and cytoplasm.

Images with 800x magnification randomly selected from each slide were used to manually count the cells at different cell cycle stages.

### Statistical analysis

SPSS statistical software (version 21.0, SPSS Inc., Chicago, Illinois, USA) was used to perform data analyses. Differences between two groups were evaluated by Student *t*-test with P < 0.05 as statistically significance. Correlation was analyzed by Pearson method.

## Results

As shown in Figure 1A, carbon tetrachloride (CCL4) induced massive hepatocyte necrosis around the central venous region (zone 3) while the periportal area (zone 1) was almost intact. Intraperitoneal injection of exogenous Tβ4 significantly reduced CCL4-induced necrosis from 36.96% (SD: 6.83%) to 29.76% (SD: 5.15%) (*P*<0.05) as regard to the area proportion (Figure 1B). These results corroborate our and others’ previous murine studies about hepatoprotective properties of Tβ4^9 10^, however so far there are no substantial *in vivo* experimental evidences about how Tβ4 influences liver regeneration. In this study effects of Tβ4 on liver regeneration were also measured through immunohistochemistry (IHC) using proliferating cell nuclear antigen (PCNA) as proliferation marker^11^. Our results (Figure 2A) demonstrated that at 48 hours after CCL4 insults most cytoplasmic PCNA-positive hepatocytes appeared around the periportal area (zone1) while less hepatocytes in the area (zone 2) adjacent to the necrosis present cytoplasmic PCNA stainings. Exogenous Tβ4 treatment prominently and obviously increases the number of PCNA-positive hepatocytes in the adjacent area of necrosis. PCNA-staining intensities were also substantially enhanced by exogenous Tβ4. Both mean nuclear and cytoplasmic H-score were significantly higher in CCL4+Tβ4 group compared with CCL4 group (Figure 2B; Nuclear H-score: 54.82±12.48 *vs.* 40.42±6.08, *P*<0.05; Cytoplasmic H-score: 72.58±9.75 *vs.* 46.48±6.88, *P*<0.05). Pearson correlation analyses showed that H-score negatively related to percent of necrosis area (Figure 3A and 3B). In order to further verify the effects of exogenous Tβ4 on hepatocytes proliferation cells at different cell cycle stages were counted. As indicated in Figure 4A, cells with deep-blue nuclear staining were at G0 stage and with light-brown nuclear staining were at G1 stage; cells with deep-brown nuclear staining were at S stage; cells presenting diffused cytoplasmic staining with or without speckled nuclear staining were at G2 stage; cells at M stage showed diffused cytoplasmic staining and deep-blue chromosomal staining^12^. Figure 4B showed that more hepatocytes at S and G2 stage (S stage: 24.36%±8.48% *vs.* 13.93%±4.21%, *P*<0.05; G2 stage: 36.80%±7.18% *vs.* 20.52% ±5.82%, *P*<0.05) and less hepatocytes at G1 stage (25.97%±5.86% *vs.* 48.28%+4.66%, *P*<0.05) were seen in CCL4+Tβ4 group compared with CCL4 group. These results suggested that intraperitoneal injection exogenous Tβ4 significantly enhanced liver regeneration at 48 hour after CCL4 insults.

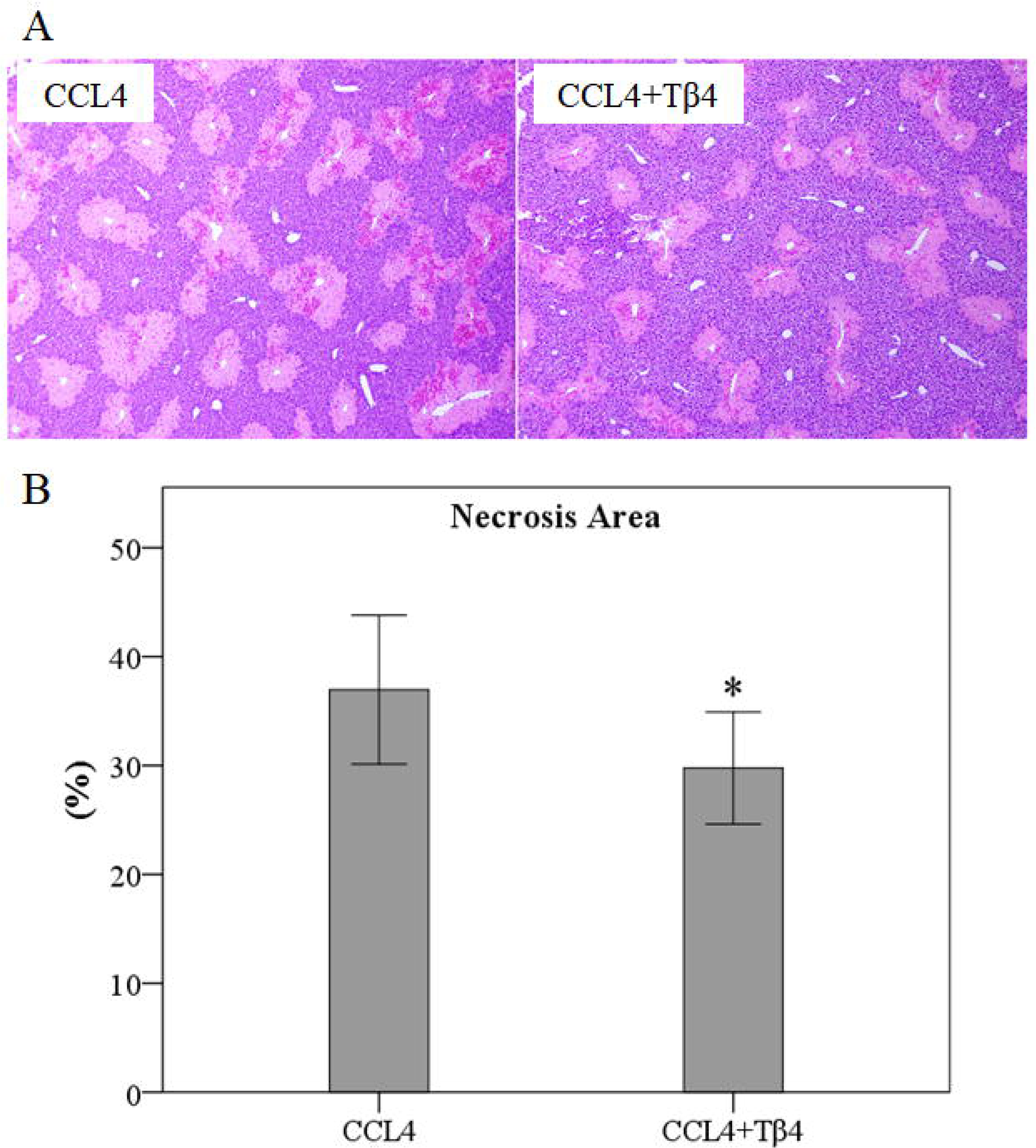

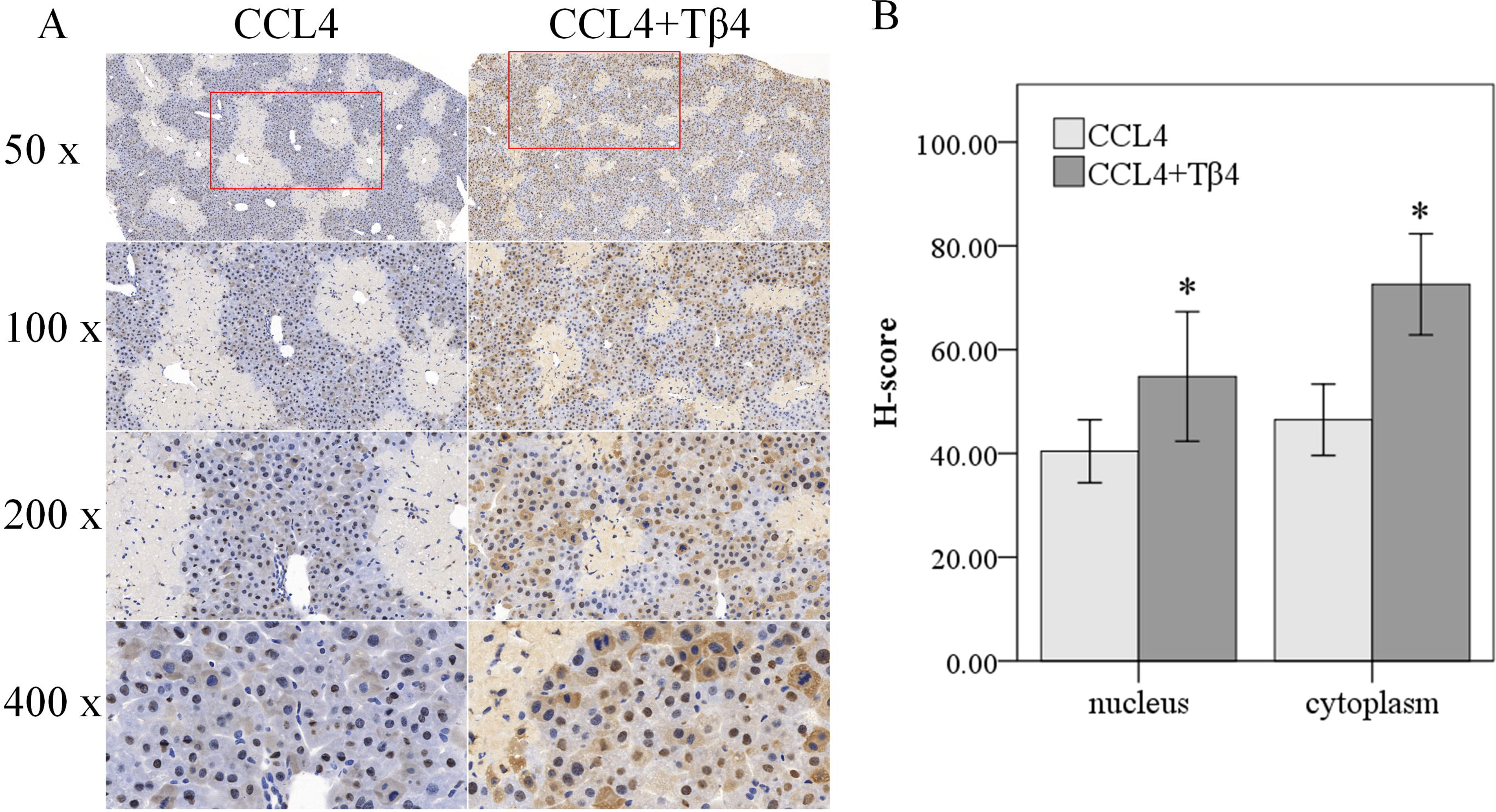

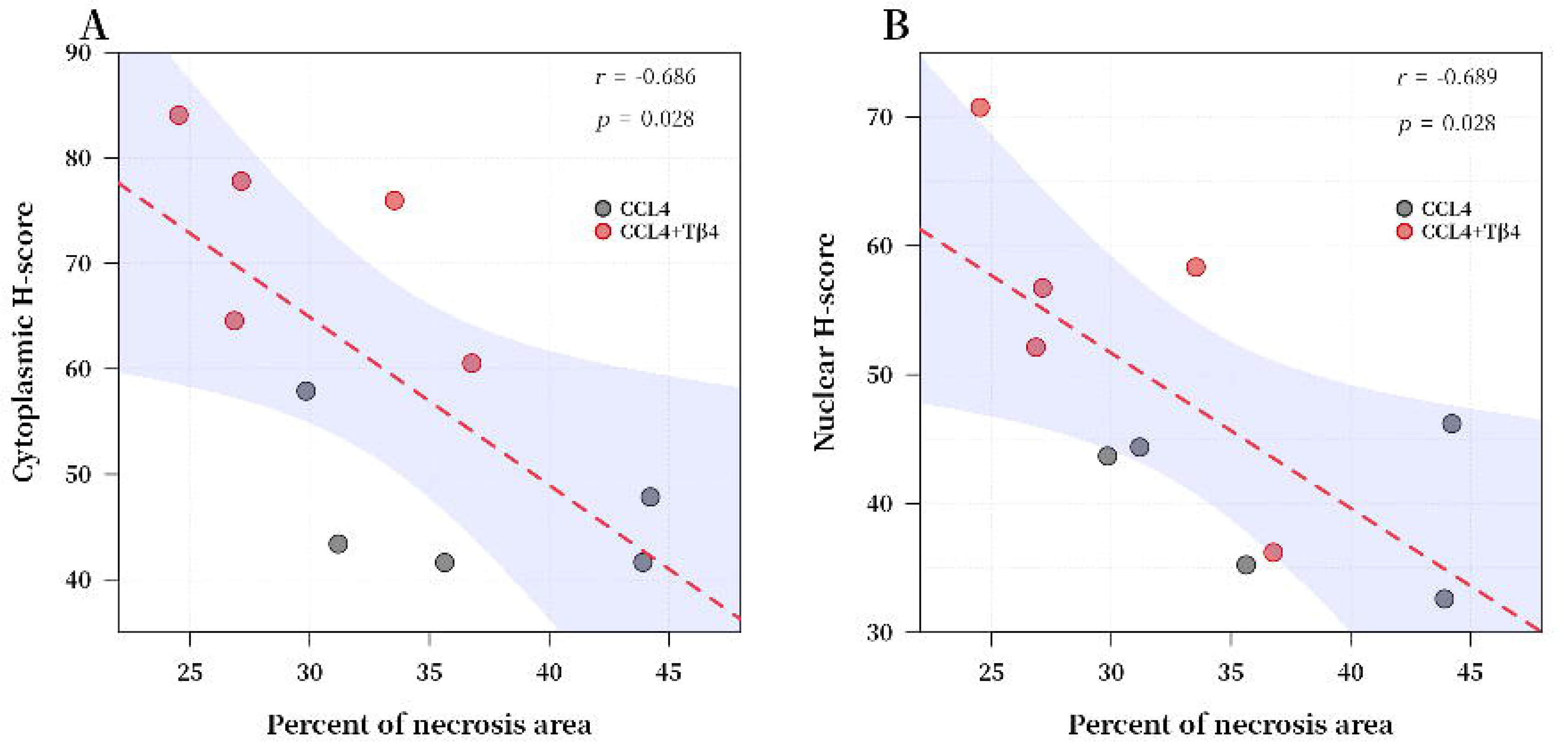

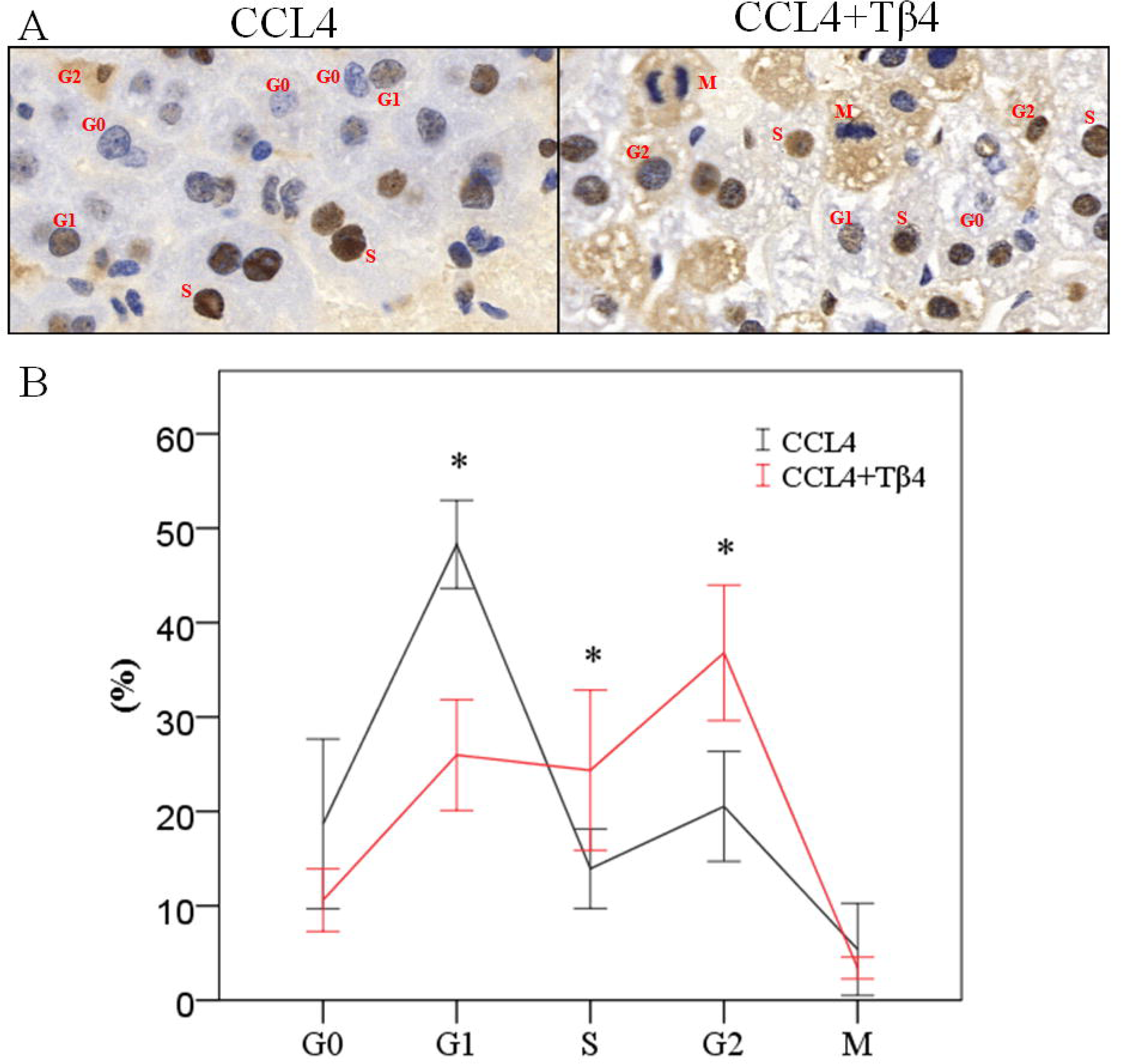

## Discussion

Clinical studies revealed that serum Tβ4 levels were significantly decreased in patients with liver failure, whereas levels markedly rebounded in those who recovered, suggesting that Tβ4 levels may reflect the process of liver injury repair^13^. Furthermore, hypermethylation of the Tβ4 promoter predicted a poor short-term prognosis in patients with acute-on-chronic hepatitis B liver failure^14 15^, and methylation levels were decreased and Tβ4 expression levels were increased by glucocorticoid therapy in patients with acute-on-chronic hepatitis B-induced liver failure^15^. Serum Tβ4 levels in patients with nonalcoholic steatohepatitis (NASH) were also significantly reduced^16^. All these clinical evidences suggested that Tβ4 played an important role in chronic liver disease.

Over the past 15 years hepatoprotective and anti-fibrosis activities of Tβ4 have attracted more researchers’ eyeballs. In 2012, Reyes-Gordillo *et al.* reported that in a CCL4-induced rat acute liver injury model exogenous Tβ4 attenuated liver injury and reduced α-SMA expression, a marker of HSC activation ^17^, which firstly point out the potential activities of exogenous Tβ4 to inhibit liver fibrosis. From then on, multiple studies from independent research teams were done to identify the role of Tβ4 in liver injury and fibrosis. Exogenous Tβ4 treatment inhibited HSC activation and proliferation and induced HSC apoptosis by suppressing AKT activation ^18 19^. Through downregulating TGF-β receptor II (TGF-βRII) ^20^, inhibiting PDGF/PDGFR and TGF-β/Smad pathways ^9 21^, the MAPK/NF-κB pathway ^22^, Notch signaling ^23^ and inducing matrix metalloproteinase (MMP) expression ^24^, exogenous Tβ4 suppressed liver fibrogenesis. Exogenous Tβ4 also acts on hepatocytes by exerting antioxidant and anti-inflammatory activities ^19^, reducing hepatocyte death and inhibiting liver fibrogenesis ^22 25^. Furthermore, exogenous Tβ4 protected liver sinusoidal endothelial cells (LSECs) from cell death, promoted LSECs proliferation and maintained LSECs fenestration, thereby contributing to the suppression of fibrosis development and progression ^26^. Although Youngmi Jung team at Pusan National University (Korea) proved endogenous Tβ4 played an active role of promoting liver fibrosis in Tβ4 gene knockout animal models^27^, at present anti-fibrosis activities of exogenous Tβ4 is an established fact by many studies.

In this study *in vivo* effects of exogenous Tβ4 on hepatocyte proliferation after CCL4 insults were investigated with PCNA as a proliferation marker^11^. Our results showed that at 48 hours after CCL4 treatment hepatocytes proliferation occurred mainly at periportal area (zone 1), while Tβ4 treatment increase the proliferative hepatocytes around the necrosis area (zone 2) and the whole levels of liver proliferation status. So far exogenous Tβ4 has not been proved to be mitogenic to hepatocytes and so the underlying molecular mechanisms are needed to further investigated in the future. Our results and conclusions are still preliminary and worthy of partial hepatectomy experiments to confirm.

Note: These data and results have been done many years ago. Because of failure of funding application and practical difficulties and constraints further investigations can’t be done possibly. I wish these data would be helpful to other researchers’ future studies.

